# Expanding Reverse Genetics of Positive-Strand RNA Viruses: Optimised Rescue Platforms and Construction of a Novel Fluorescent Reporter Nidovirus

**DOI:** 10.64898/2026.08.25.746995

**Authors:** James R. Potter, Helen Mostafavi, Alberto A. Amarilla, Ryan A. Johnston, Rhys H. Parry, Margus Varjak, Alain Kohl, Alexander A. Khromykh, Natalee D. Newton, Jody Hobson-Peters

## Abstract

Reverse genetics systems are crucial for facilitating the precise manipulation of viruses across a wide spectrum of translational and fundamental research pipelines. Here, we compared Circular polymerase extension reaction (CPER), Gibson assembly, and infectious subgenomic amplicons (ISA) for bacteria-free recovery of a positive sense RNA virus. Through optimisation of CPER, we demonstrated accelerated virus recovery and enhanced viral yields. We further investigated strategies to improve rescue efficiency across diverse positive-sense RNA virus families through incorporation of alternative promoters and non-coding elements. To evaluate the performance of the *Aedes aegypti* polyubiquitin promoter (AePUb) in tandem with a hammerhead ribozyme (HH Rbz) and a polymerase pause site for virus recovery in insect cells, we constructed a new fluorescent reporter genome using a 20 kb insect-specific mesonivirus. In vitro recovery by CPER of the mesonivirus was achievable in 1 day when using AePUb with HH Rbz, in comparison to a four-day recovery when using the minimal OpIE2-CA promoter. These elements were additionally assessed for rescue of the orthoflaviviruses, Binjari virus (BinJV) and dengue virus 2 (DENV-2), in insect cells (using AePUb); or in mammalian cells (using the CMV promoter) and for launch of DENV2 and SARS-CoV-2. Both BinJV and DENV-2 demonstrated improved rescue with the AePUb promoter and HH Rbz. However, the addition of the HH Rbz and the polymerase pause site to the CMV linker fragment showed no substantial differences to the standard CMV promoter systems for both DENV-2 and SARS-CoV-2, highlighting the context-specific benefits of their implementation. In summary, we demonstrated that a potent constitutive promoter system and a hammerhead ribozyme enhance the efficiency of positive-sense RNA virus rescue using CPER.

**Importance:** Reverse genetics systems are often limited by plasmid instability and variable efficiency of promoters across diverse cell lines. Extensive comparative approaches have yielded improvements across a variety of systems, however, there has been a paucity of publications that empirically compare novel advancements to established approaches. Here, we formalised and compared a series of reverse genetics advancements in the form of bacteria-free assembly methods, host promoters, pause sites, and ribozymes. These streamlined approaches expedite the existing methodologies and provide fundamental improvements to the field of synthetic virology. The advancements herein may support applications requiring efficient recovery of low fitness mutants and diverse mutational libraries and barcoded virus populations.

## Introduction

Reverse genetics systems enable researchers to genetically manipulate viruses to investigate viral function, produce attenuated clones, and engineer recombinant expression vectors^1–3^. Historically, reverse genetics systems of viruses have been reliant on the recombinant cloning of viral cDNA into plasmids or equivalent vectors (e.g. BACmids, COSmids, Yeast vectors) using restriction endonucleases or assembly reactions^4^. However, a limitation of these classical cloning systems is that propagating cDNA through bacteria introduces a selection step in which mutant cDNA clones can outcompete the desired virus cDNA clone^5–7^. This problem is exacerbated for viral sequences, which are often toxic in bacteria^8^. The bacterial shuttling process also requires a backbone with expression cassettes in the vector which can be detrimental to downstream expression efficiency upon transfection^7,9^.

The first bacteria-free viral reverse genetics system to circumvent these issues, Circular Polymerase Extension Reaction (CPER), was implemented for the construction of an infectious clone of the orthoflavivirus, West Nile virus in 2013^10^. Since then, a variety of other bacteria-free methods such as Gibson assembly and Infectious Subgenomic Amplicons (ISA) have been developed for diverse families of RNA viruses^11–14^. These methods rely on viral cDNA PCR amplification of dsDNA fragments containing homologous overhangs to adjacent fragments. These fragments comprise both the entire viral genome and an additional dsDNA fragment cassette incorporating a 5’ end host-specific promoter and a 3’ end hepatitis delta virus ribozyme (HDVr) to ensure viral 3’UTR authenticity by cis-acting ribozyme self-cleavage. The assembly results in a circular dsDNA topology for Gibson assembly and CPER, while ISA is designed to result in a linear product (Fig. 1B)^8^. Overall, these advancements have led to the widespread adoption of bacteria-free systems for both molecular virology studies and for manufacturing processes which use virus vectors and have regulatory constraints on bacterial-based production due to potential endotoxins, such as the manufacture of vaccines and therapeutics^11,15–20^. The use of CPER has expanded considerably within the past decade and has been applied to reverse genetics of positive-sense RNA viruses of numerous families, including mosquito-and tick-borne orthoflaviviruses, insect-specific flaviviruses, alphaviruses, nidoviruses, including SARS-CoV-2 and caliciviruses^16,21–25^. For dual-host viruses, such as the orthoflaviruses and alphaviruses, CPER has facilitated virus recovery in mammalian or insect cell cultures^16,22,26^. More recently, CPER has also been adapted for negative-sense RNA virus systems, with studies demonstrating its utility for mononegaviruses, including respiratory syncytial virus and rabies virus ^27,28^. Collectively, the application of CPER and related methodologies has had a substantial impact on the study of a diverse range of viruses.

**Figure 1.**
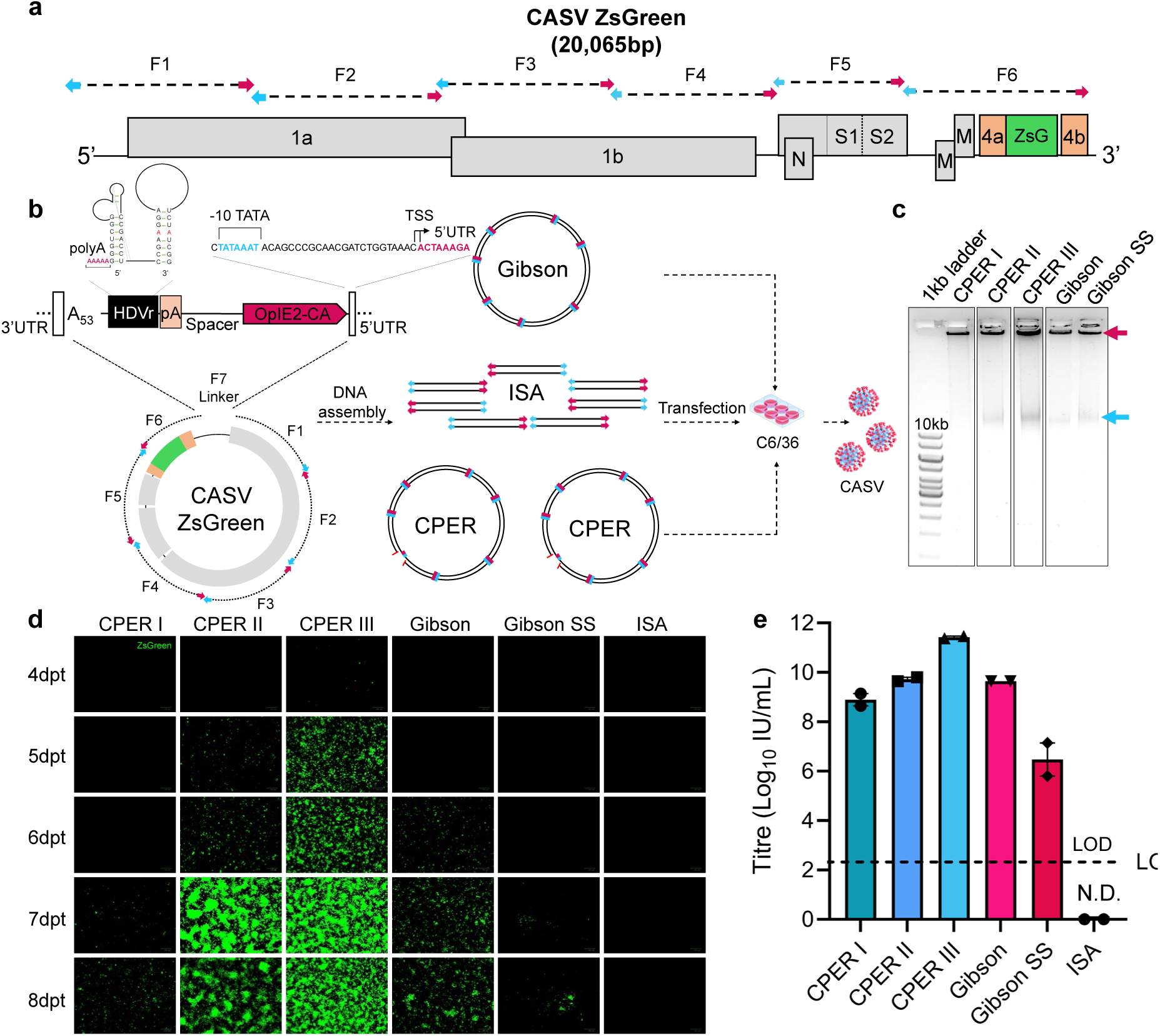
Reverse genetics methodologies comparison for the recovery of a CASV reporter virus. **A**) Schematic of CASV-ZsGreen reverse genetics construct comprised of the CASV genomic fragments with overlapping ends, ZsGreen reporter sequence (ZsG) encoded downstream of the putative ORF4a and separated by a FMDV 2A ribosomal skipping peptide **B**) The overlapping genomic fragments are circularised with a linker fragment containing the last 35 nucleotides of the CASV 3’UTR, 53 A’s, hepatitis delta virus ribozyme (HDVr), SV40 pA signal for transcription termination, spacer sequence, OpIE2-CA promoter and the first 25 nucleotides of the CASV 5’UTR. **C**) 0.8% agarose gel of 1/10 volume of the total assembly reaction for each methodology. ISA was excluded as it is not an assembled product prior to transfection. The expected band size for each reaction is annotated with a blue arrow (20 kb). Concatemeric or branched DNA formation in wells are highlighted with a red arrow. CPER I 20 cycles, extension time 15 minutes; CPER II 35 cycles, extension time 15 minutes; CPER III 20 cycles, extension time 7 minutes. SS = addition of single-stranded DNA binding protein. **D**) Fluorescence imaging using a ZOE Cell Imager (Bio-Rad) of C6/36 cell monolayers post-transfection with CASV-ZsGreen CPER, Gibson assembly and ISA products. Monolayer imaging was performed daily from 4 days post-transfection. **E**) Mean titres of CASV-ZsGreen virus harvested from transfected wells at 8 days post-transfection. Two biological replicates are shown. LOD = Limit of Detection = 2.3 log_10_IU (infectious units)/mL. Error bars = Standard error of the mean (SEM).

In addition to the increasing adoption of CPER and other bacteria-free reverse genetics systems for virus recovery, there have been advancements focused on the optimisation of transcription efficiency and transcript processing. These investigations have been primarily targeted towards the reverse genetics system promoter elements and associated ribozymes in the non-coding region of constructs (typically referred to as a “linker” in the literature)^8,16,29,30^. In the context of promoters, mammalian reverse genetics systems have used a CMV immediate-early promoter recognized by RNA polymerase II (RNA Pol II) or an RNA polymerase I promoter to avoid steric hindrance of the capping mechanism in viruses that use IRES-mediated translation^31^. To enhance efficiency, some reverse genetics systems for bunyaviruses and paramyxoviruses have used a more efficient T7 bacteriophage promoter in a BSR-T7 (baby hamster kidney) cell line which expresses a highly processive T7 polymerase^32,33^.

Insect cell reverse genetics systems typically employ the RNA Pol II promoter OpIE2 from baculovirus, which was modified with a transcription start site truncation to improve its efficiency for RNA virus recovery^30,34^. However, for reverse genetics systems tailored to mosquito cells, alternative promoters with substantially stronger activity have been identified. Among these, the *Aedes aeqypti* polyubiquitin promoter (AePUb) has been shown to drive robust gene expression in vivo and in vitro^35–37^, yet its application in mosquito cell-based virus reverse genetics systems remains largely unexplored. Recent studies have also employed a polymerase pause site sequence derived from the human α2 globin gene^29^. This sequence was inserted in the spacer region (upstream of the promoter sequence) for CPER-based launch of SARS-CoV-2 to minimise RNA Pol II read-through in the linker region. Its inclusion was associated with enhanced virus recovery efficiency, albeit in conjunction with the alteration of other cell line variables and additional methodological modifications, therefore making the contribution of the pause site difficult to isolate^29^. Another important consideration in reverse genetics systems is the effect of promoter truncation on transcriptional efficiency while still ensuring generation of a native viral 5′ UTR terminus without the addition of extraneous residues. Precise trimming of negative-sense viral genomes, including Nipah virus and bunyaviruses, has been achieved through incorporation of a hammerhead ribozyme sequence (HH Rbz) immediately upstream of the viral antigenome 5′ terminus^38,39^.

Collectively, while advancements in bacteria-free reverse genetics systems have greatly expedited workflows by obviating common methodological bottlenecks, implementation of these systems remains technically demanding and frequently results in experimental failure, particularly for less optimised insect virus-based systems. Here, we report a systematic analysis of multiple bacteria-free reverse genetics approaches, with a particular focus on CPER, and empirically define features that enhance virus recovery in both mammalian and mosquito cells. Importantly, our optimisations enabled recovery of a large-genome mosquito virus in mosquito cells in 24 hours. In addition, several modifications to our own procedures were incorporated to provide practical insights and a useful reference point for synthetic virologists seeking to employ reverse genetics systems in either insect or mammalian cell systems.

## Results

### Comparison of bacteria-free reverse genetics methods for the rapid synthesis of RNA viruses

First, we compared CPER, Gibson assembly, and ISA for their respective efficiencies in recovery of positive-sense single-stranded RNA (+ssRNA) viruses using a large mesonivirus (nidovirus), Casuarina virus (CASV), as a representative template^16,40^. We have previously reported the construction of a CASV infectious clone using CPER to produce an authentic recapitulation of the wild-type isolate in vitro^16^. However, to enable comparisons of virus recovery across the reverse genetics systems in real-time, a novel fluorescent reporter construct was designed in which a ZsGreen-coding sequence was placed downstream of the putative CASV accessory ORF4a (Fig 1A). To partition expression of the ZsGreen cassette from the upstream ORF4a protein, a foot-and-mouth-disease virus ribosomal skipping peptide (FMDV 2A)^41,42^ was encoded at the C-terminal end of ORF4a (Fig 1A). Six contiguous genomic fragments were amplified with 20-40bp homologous overhangs and were generated from CASV cDNA template or a gBlock for the reporter cassette (Figs 1A, B). The 5’ and 3’ genomic ends were designed to be joined by a linker fragment consisting of a 3’ end hepatitis delta virus ribozyme (HDVr) followed by a simian virus 40 (SV40) late polyA motif (pA), a spacer region, and an optimised version of an immediate-early promoter derived from Orgyia pseudotsugata multicapsid nuclear polyhedrosis virus (OpMNPV), termed OpIE2-CA^30^.

For CPER assembly, three variations of cycling conditions were trialled in which the number of cycles and the extension time were varied. CPER I was conducted with 20 cycles and an extension time of 15 minutes and represented the original protocol used to launch the CASV infectious clone^16^. CPER II was designed to trial an increase to 35 cycles, while CPER III had a reduced extension time of 7 minutes (1 min per kb of the longest fragment) and maintained a cycle number of 20. The Gibson assembly was incubated at 50 °C for 4 hours. Variations for cycling conditions were not included as Gibson assembly is an isothermal method. The assembly reaction for Gibson was also optionally supplemented with a single-stranded DNA binding protein (SSBP) under the rationale that it prevents potential endonuclease activity and can increase polymerase processivity under extended exposure to high temperatures^43^. In contrast to CPER and Gibson protocols, ISA requires no assembly steps prior to transfection.

For all protocols, the same amplified PCR fragments encoding the full-length CASV reporter virus were used to ensure that differences in DNA quality did not confound comparisons of assembly efficiency. Following assembly for CPER and Gibson protocols, the resulting product was assessed by gel electrophoresis (Fig 1C). All methods demonstrated substantial banding in the gel wells, which is consistent with concatemeric or branched DNA formation which occurs during assembly ^16^. The CPER III, CPER II, Gibson, and Gibson with SSBP methods showed prominent band formation above 10kb (assembled product is 20kb), indicative of successful assembly. The CPER I displayed no banding, consistent with reduced assembly (Fig 1C).

The assembly products were transfected using TransIT-LT1 into *Aedes albopictus* C6/36 cells for virus recovery. Fluorescence was assessed in daily intervals and culture supernatants were harvested at 8 days post transfection (dpt) and titrated onto C6/36 cells before assessment of infectious virus titres (Fig 1D, E). The CPER III protocol with 20 cycles and a 7-minute extension time resulted in both the earliest detectable fluorescence (4 days, as sparse visible fluorescence) and the highest infectious virus titre at 8 dpt, which were greater than all other conditions, including Gibson assembly (Fig 1E). CPER III conditions recovered virus three days earlier than the original protocol (CPER I), while increasing the cycle number (CPER II conditions) also resulted in reporter virus recovery two days faster than the original conditions (Fig 1D). This comparative benefit shows the importance of cycling conditions for the efficiency of CPER assembly and emphasises the need to assess CPER assemblies by gel electrophoresis as a proxy for efficiency prior to transfection.

Fluorescent imaging indicated that ISA was the only method that resulted in no detectable expression, which was further supported by the absence of infectious virus in titration assays (Fig 1D, E). Addition of SSBP to Gibson assembly had a deleterious effect, resulting in both slower rescue and lower yields of virus. This may be due to SSBP steric hindrance of polymerases, ligases, and exonucleases during assembly.

### An Aedes aegypti promoter enhances insect-based reverse genetics of CASV in insect cells

Studies of arboviruses and the development of insect-derived vaccines are dependent on robust reverse genetics systems for mosquito cell lines^24,25,44^. However, there has been minimal focus in the literature on enhancing the recovery efficiency of these systems. Here, the optimised CPER III protocol was implemented in a comparative reverse genetics study of different promoter sequences in the C6/36 mosquito cell line for rescue efficiency. Promoter sequences are key determinants of viral transcript production upon transfection into the cell nucleus and are therefore one of the key parameters for manipulation in different host-cell contexts. The minimal OpIE2-CA promoter has been used broadly across insect-cell reverse genetics systems despite not being a mosquito-derived promoter^16,24,25,30,45^. To improve this existing approach, the AePUb promoter^35–37^ was implemented here and assessed for its comparative recovery efficiency.

The AePUb promoter was assessed in parallel with the full-length OpIE2 and the optimised truncated OpIE2-CA promoter for efficiency of virus rescue by insertion into the linker fragment in the CASV-ZsGreen construct (Fig 2A). The AePUb promoter sequence has a mapped transcription start site 826bp upstream of the utilised promoter sequence^35^. To maintain the transcription-enhancing elements of the promoter downstream of the transcription start site, a hammerhead ribozyme sequence (HH Rbz) was placed immediately after the promoter sequence (termed AePUb HH, Fig 2B). The HH Rbz autocatalytically cleaves away from the transcript produced downstream of it (the viral 5’UTR, Fig 2C) and therefore prevents the unnecessary and potentially inhibitory addition of nucleotides to the viral transcripts, which is vital in lieu of a transcription start site truncation^46,47^.

**Figure 2.**
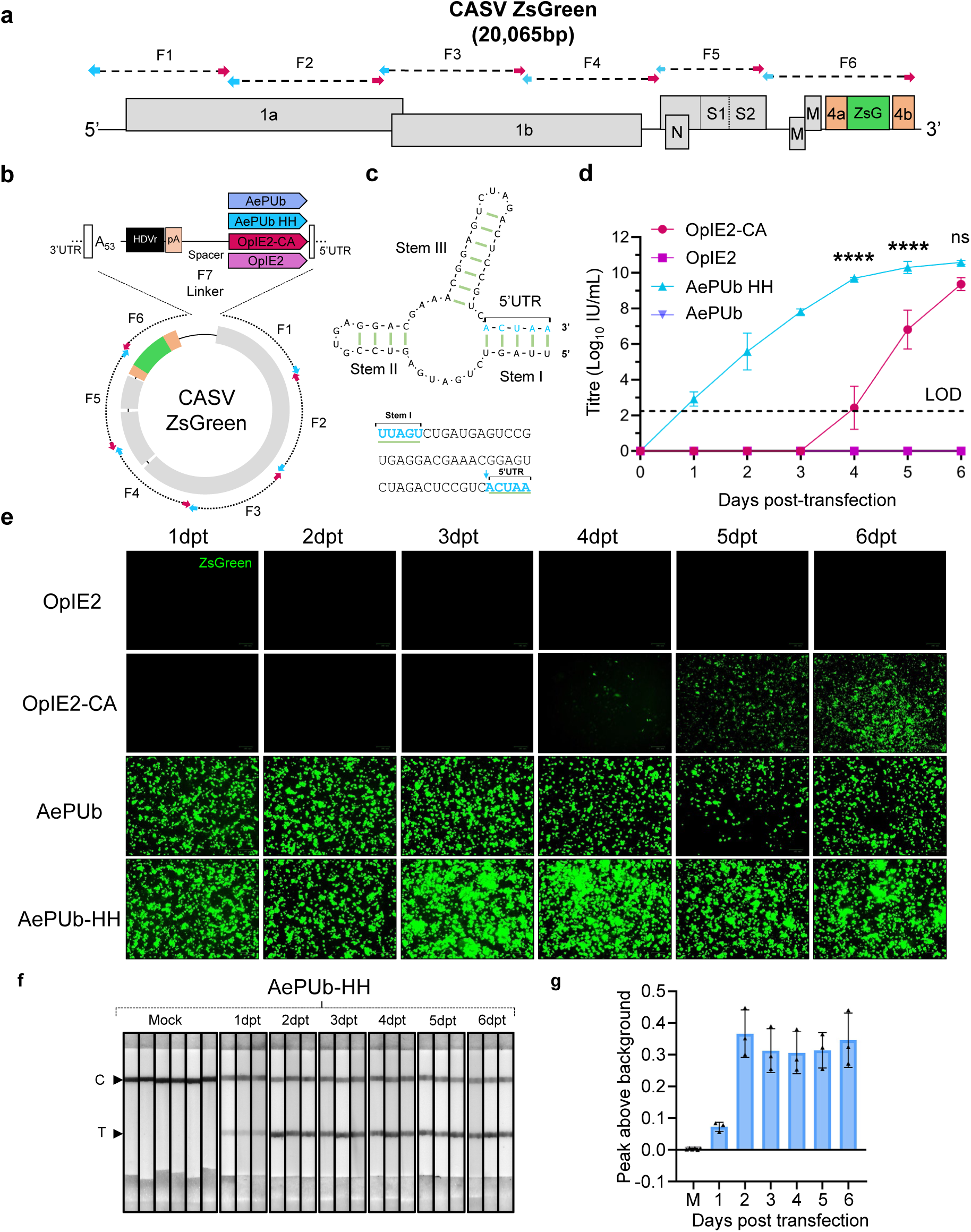
Comparison of promoter systems for CASV-ZsGreen reverse genetics optimisation. **A)** Conceptual schematic of the CASV-ZsGreen genome and the overlapping fragments for the CPER assembly. F1-F6 represent each separate fragment, with the arrows corresponding to the homologous overlaps of adjacent fragments. **B)** The overlapping genomic fragments were circularised with a linker fragment containing the last 35 nucleotides of the CASV 3’UTR, 53 A’s, hepatitis delta virus ribozyme (HDVr), SV40 pA signal for transcription termination, spacer sequence and one of four different promoters: full length AePUb, AePUb with HH Rbz (AePUb-HH), OpIE2-CA, OpIE2 (full length). **C)** Predicted secondary structure of the HH ribozyme and the first five nucleotides of the CASV 5′UTR generated using RNAFold and visualised with VARNA. **D)** Growth kinetics of CASV-ZsGreen for each transfection in 24-hour time points assessed by TCID_50_ in C6/36 cells using a ZsGreen readout. Experiment was performed in 3 biological replicates. Statistical significance of the differences in mean titres between different promoters at each timepoint was determined by two-way ANOVA using Tukey’s multiple comparisons and is indicated (**** p<0.0001, ns = not significant p>0.05). LOD = Limit of Detection = 2.3 log_10_ IU/mL. Error bars = SEM. **E)** Fluorescent imaging of transfected C6/36 cells at daily intervals between one dpt and six dpt. **F)** Lateral flow test detection of AePUb-HH-driven CASV-ZsGreen in culture supernatants (assessed in panel D) taken daily post-CPER transfection using anti-CASV S monoclonal antibodies (mAbs 5D3 and 9D7). **G**) Lateral flow test line peak above background of triplicate samples in **F** as analysed using the Leelu reader. Error bars = SD. Panel F shows images of the corresponding LFT strips. M = mock supernatant, D1 – D6 = the day post-transfection, C = control line, and T = test line indicating binding of virus by mAb 9D7.

To ensure accurate comparison between promoter sequences and minimise variation arising from differences in nucleotide composition and thus melting temperatures of the junction between the linker region and the first fragment of the CASV genome, the promoter-containing linker fragment was first fused to genomic fragment 1 (F1) by overlap extension PCR prior to the CPER assembly with the remaining five CASV-ZsGreen genomic fragments. Following transfection into C6/36 cells, both AePUb and AePUb-HH-driven CPER conditions showed substantial fluorescence at just 24 hours post-transfection (Fig 2E). The full-length OpIE2 CPER transfected cells showed no fluorescence at any time-point, while the truncated OpIE2-CA CPER showed fluorescence at 4 days post-transfection, which was consistent with the results obtained during the protocol optimisation (Fig 1).

Supernatant from all four CPERs comparing the different promoters was harvested and titrated onto C6/36 cells for assessment of the rescue kinetics (Fig 2D). The AePUb-HH promoter produced the highest titre of virus at all time points, peaking at ∼10^10^ IU/mL, and that viable virus was detectable by 1-day post-transfection. This was both faster than the original OpIE2-CA promoter, which produced detectable infectious virus at 4 days post-transfection, and produced significantly more virus at days 4 and 5 post-transfection. Despite the AePUb promoter (without HH Rbz) CPER-transfected cells showing strong fluorescence at 24 hours post-transfection (Fig 2E), upon passaging there was no infectious virus present above the limit of detection (Fig 2D). This suggested an inhibitory effect of the additional nucleotides on the viral transcripts produced from the AePUb promoter sequence and reinforces the requirement of the HH Rbz in this context. One potential explanation is that in the absence of the HH Rbz self-cleavage, there is an excess of nucleotides appended to the 5’UTR, which in turn interfere with its downstream functions related to packaging and dsRNA replication. This is distinct from the impact on initial transcripts produced from the promoter, which are still capable of expressing viable protein, as exemplified by the strong ZsGreen expression.

As promoter-driven fluorescence confounded discrimination from virus replication-derived fluorescence in the AePUb transfections, interpretation of daily fluorescence imaging was unreliable. To provide a rapid independent measure of virus rescue and corroborate quantitative supernatant titrations, a lateral flow test (LFT) for CASV virion detection was assessed using the anti-CASV S monoclonal antibodies 5D3 and 9D7^48^. The LFT results were consistent with the titration data obtained from AePUb-HH CPER transfections and confirmed successful rescue of CASV-ZsGreen within 1-day post-transfection, with a clear positive test line detected in culture supernatants (Fig 2F, G). Peak test line intensity was observed from day 2 post-transfection onwards. Collectively, these findings demonstrate that the LFT provides a rapid and effective alternative for assessing virus rescue.

Recent work by Liu and Gack (2023) in the context of SARS-CoV-2 reverse genetics employed a polymerase II pause site sequence derived from the human α2 globin gene in the linker sequence^29^. This was introduced following the rationale that rescue could be enhanced by preventing the production of heterologous transcripts and ameliorating any potential interference with polymerase recruitment to the promoter ^29^. Following this same rationale, the pause site was integrated into the linker sequence with AePUb-HH as a component of the CASV-ZsGreen reverse genetics system established here (Fig 3A, B). This was compared for efficiency of CASV-ZsGreen synthesis in the absence of the pause site. ZsGreen fluorescence and virus rescue kinetics assessment post-transfection revealed no significant difference between the two linkers in terms of recovered virus titre, nor to the timepoint for virus rescue (p>0.05, Fig 3C, D).

**Figure 3.**
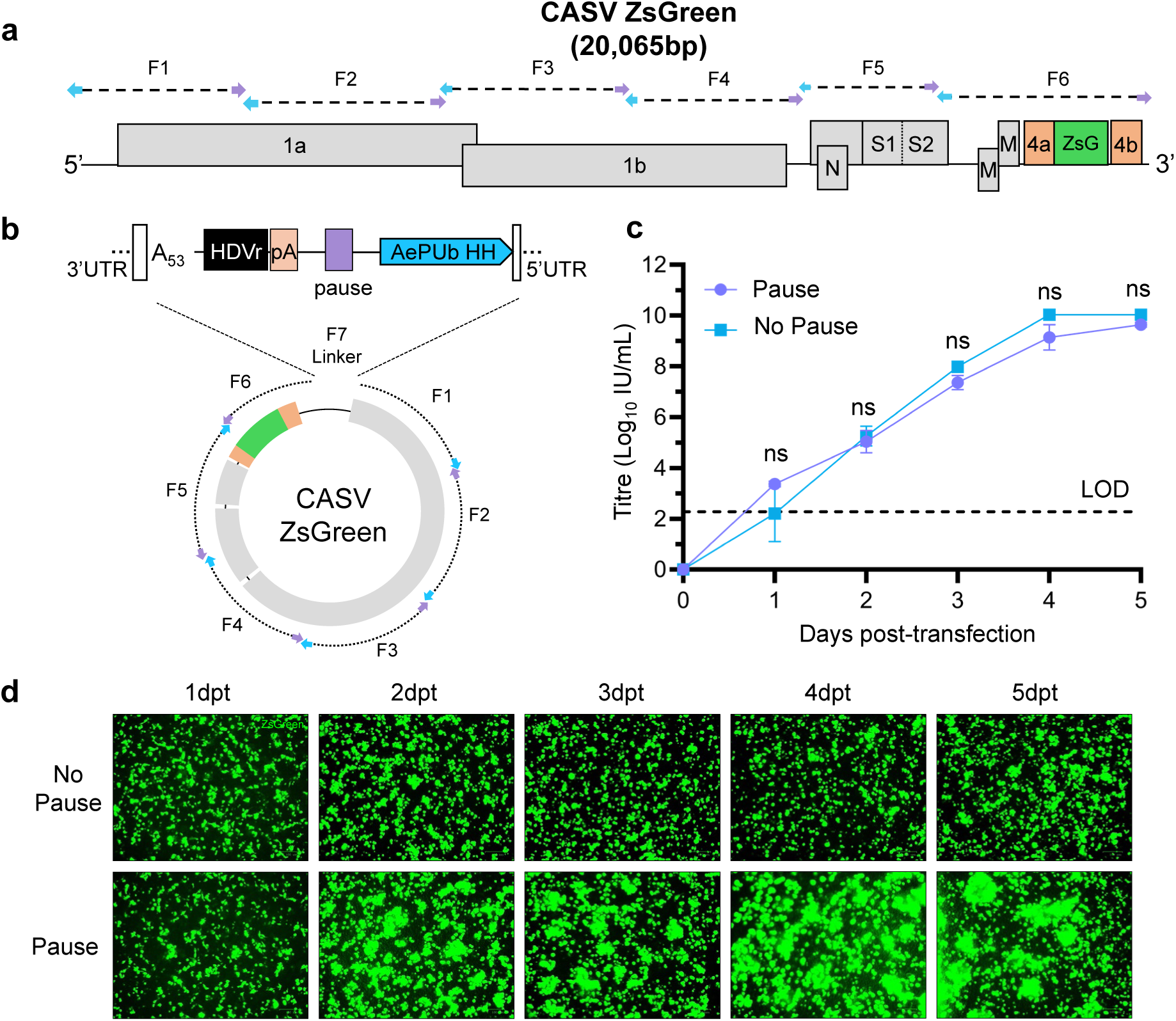
Recovery kinetics of CASV ZsGreen using a polymerase pause site. **A)** Schematic of the CASV genomic fragments used for assessment of virus recovery efficiency upon inclusion of a polymerase pause sequence, which was optionally included into the linker fragment upstream of the AePUb-HH promoter **B)** The overlapping fragments of CASV were circularised by CPER with a linker optionally containing the pause site sequence shown and then transfected into C6/36 cells. **C)** Growth kinetics of CASV-ZsGreen for each transfection in 24-hour time points assessed by TCID_50_ in C6/36 cells using a ZsGreen readout. Experiment was performed in 3 biological replicates. Statistical significance of the differences in mean titres between treatments at each timepoint was determined by two-way ANOVA using Šídák’s multiple comparisons and is indicated (ns = not significant p>0.05). LOD = Limit of Detection = 2.3 log_10_ IU/mL. Dpt = days post transfection. Error bars = SEM. **D)** Fluorescent imaging of CASV ZsGreen CPER-transfected C6/36 cells at daily intervals between one day post-transfection and six days post-transfection using a ZOE fluorescent microscope (Bio-Rad), with one representative well shown for each.

### Assessment of the AePUb-HH promoter system for launch of insect-specific flaviviruses

To assess the utility of the AePUb-HH promoter sequence in reverse genetics systems for other virus families, the synthesis of a fluorescent mCherry-expressing insect-specific flavivirus (ISF), Binjari virus (BinJV, BinJVmCherry), was assessed using an established design ^45^ (Fig 4A). This design positioned the first 51 amino acids of the capsid gene upstream of the mCherry cassette to preserve the cis-acting RNA elements, while the downstream full-length capsid gene was codon-optimised to prevent recombination and reporter gene loss^49,50^. BinJV has rapidly been established as a robust platform to enable orthoflavivirus structure elucidation and for the development of vaccine and diagnostic antigens^24,51,52^. Thus, it provided a prototypical example of a positive-sense RNA virus with a smaller genome than the mesoniviruses that would benefit in both basic and translational pipelines from a more efficient reverse genetics system. Again, the OpIE2 and AePUb-HH linker fragments were independently joined by overlap-extension PCR to the viral fragment 1 to circumvent the variable annealing temperatures for homologous overhangs (Fig 4B). The rescue kinetics for BinJV_mCherry_ demonstrated a comparative advantage of the AePUb-HH promoter system in comparison to using the OpIE2-CA promoter (Fig 4D, E). When using AePUb-HH, virus was rescued at day 2, while when using the OpIE2-CA promoter, virus was not detected until day 4, with these observations consistent upon assessment for replicating virus in a TCID_50_ assay and by fluorescent imaging of cell monolayers (Fig 4D, E). The AePUb-HH linker also promoted the recovery of virus at substantially higher titres at days 4 and 5, reaching a total of ∼8 log_10_TCID_50_ units compared to ∼6 log_10_TCID_50_ units for the OpIE2-CA-launched virus. As these data were in alignment with that observed for the CASV ZsGreen construct, we concluded that the AePUb-HH linker provides a substantive improvement upon existing insect-based reverse genetics systems.

**Figure 4.**
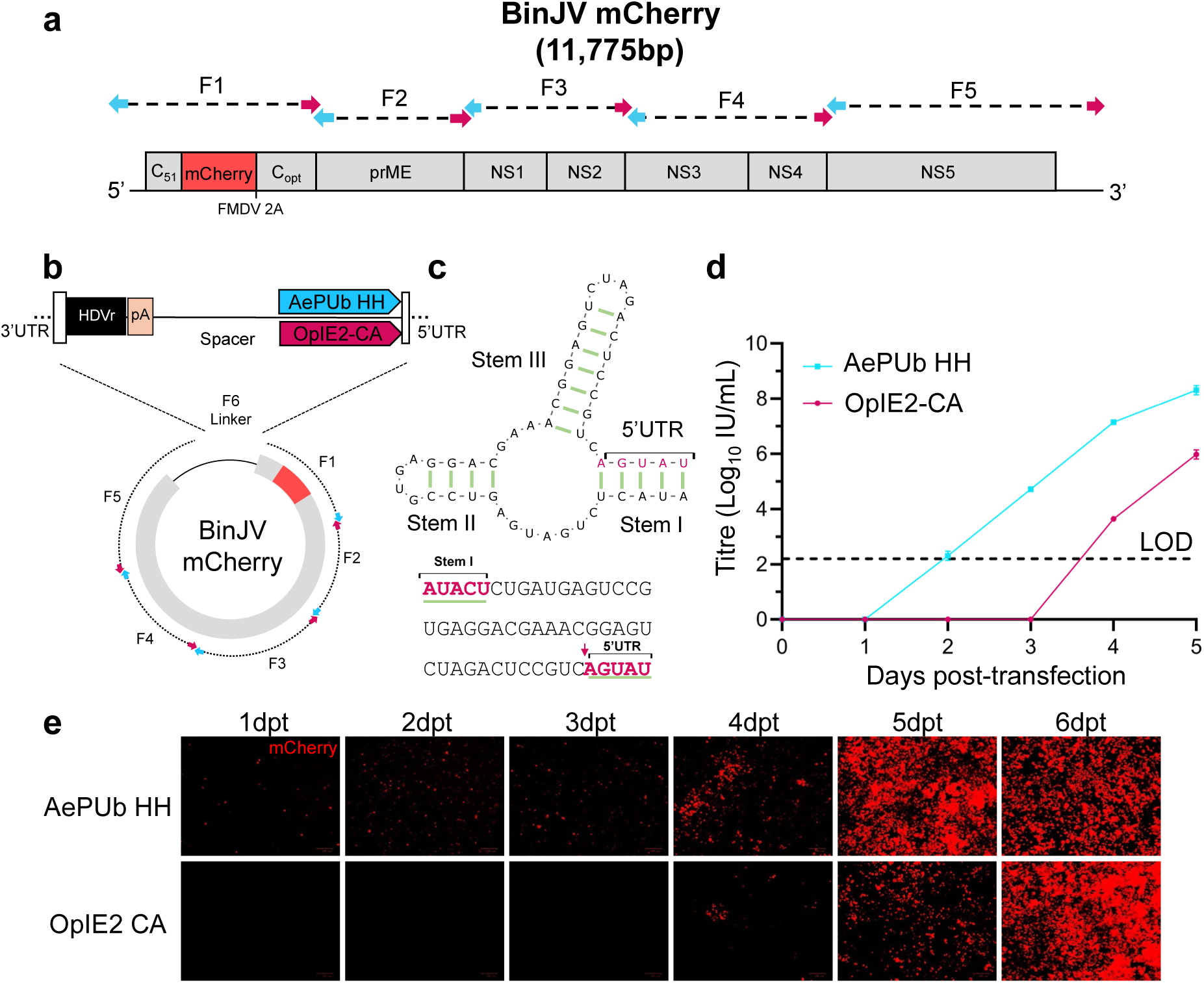
Comparison of promoter systems for BinJV_mCherry_ reverse genetics. **A**) Schematic of the BinJV_mCherry_ fluorescent virus genome and overlapping fragments. **B**) The DNA linker was incorporated by CPER following overlap extension PCR with the BinJV F1 genomic fragment and contained with either the AePUb-HH or OpIE2-CA promoter. **C**) Predicted secondary structure of the HH ribozyme and the first five nucleotides of the BinJV 5′UTR generated using RNAFold and visualised with VARNA. **D**) Kinetics of BinJV_mCherry_ viral titres, as determined by TCID_50_ using mCherry fluorescence, when harvested at daily intervals from 1 day to 5 dpt. Experiments were performed in 2 biological replicates. Error bars = SEM. LOD = Limit of Detection = 2.3 log_10_ IU/mL. **E**) Fluorescent imaging of BinJV_mCherry_ CPER-transfected C6/36 monolayers using a ZOE microscope (Bio-Rad) at daily intervals.

### Assessment of promoter and ribozyme elements for rapid RNA virus rescue in mammalian cells via CPER

Following the modifications for enhanced virus recovery in insect cells, we sought to determine if similar modifications to well-established mammalian reverse genetics systems could accelerate virus recovery. For these comparisons, dengue virus serotype 2 (DENV-2) was selected, allowing comparison of insect-and mammalian-cell launch systems using the same virus.

In previous studies, mammalian-based reverse genetics systems for DENV-2 and other orthoflaviviruses have used a minimal CMV promoter ^10,22,53^. Despite CMV’s extensive use, it has not had its transcription start site mapped experimentally in existing published studies. Therefore, the HH Rbz was appended to the CMV promoter sequence to prevent the addition of extra nucleotides to the 5’UTR of the virus transcripts (Fig 5A-C). This HH Rbz modification was coupled with the polymerase pause site, noting that the latter was shown to enhance SARS-CoV-2 rescue ^29^. While the pause site had no discernible impact on CASV rescue efficiency in insect cells, it was included here to assess its potential impact in a mammalian system.

**Figure 5.**
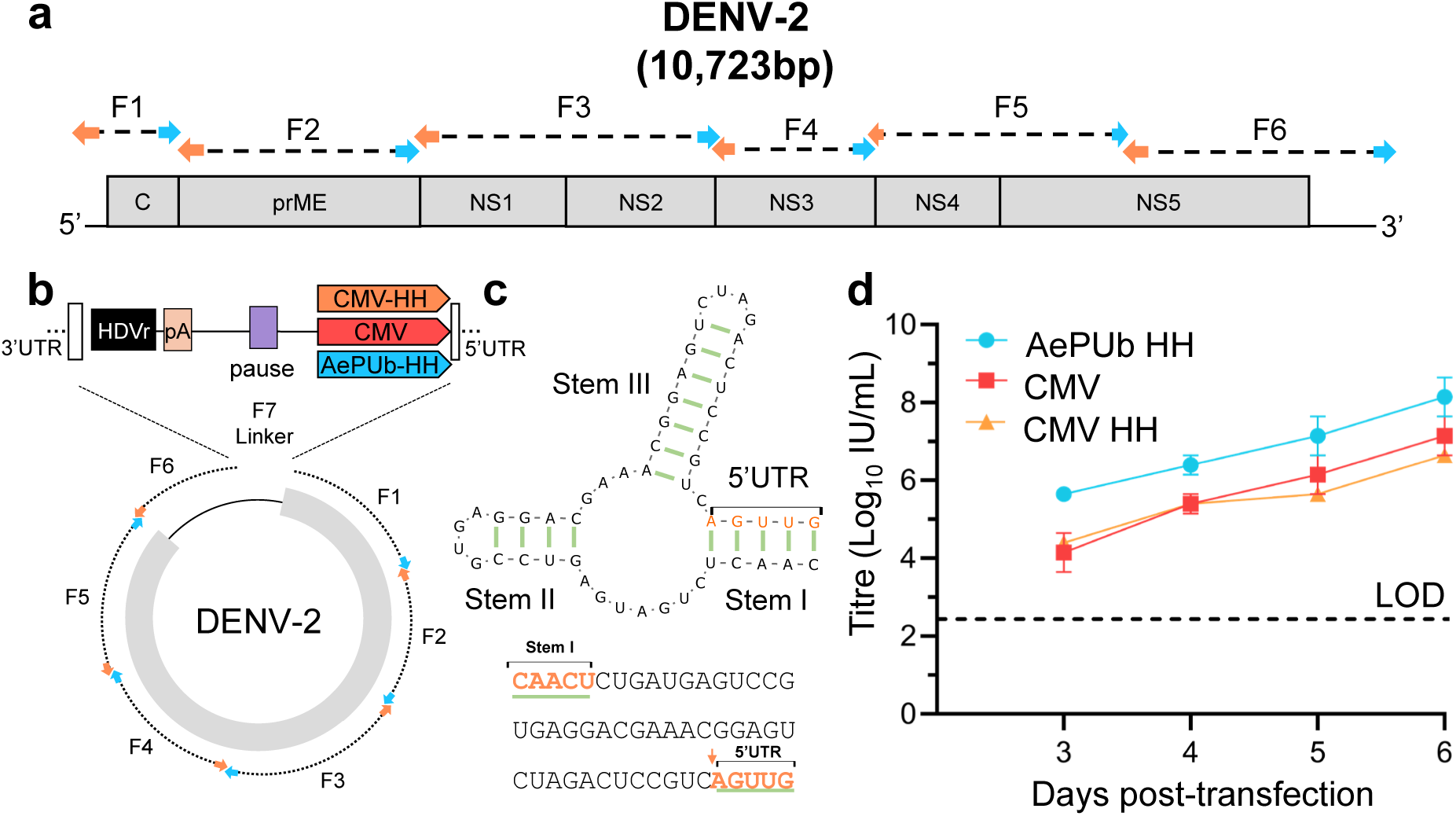
Comparison of promoter systems for DENV-2 reverse genetics. **A**) Schematic of the DENV-2 genome and overlapping fragments designed for CPER. **B**) The DNA linker was incorporated by CPER following overlap extension PCR with the DENV F1 genomic fragment and contained either the following promoter and other sequence elements: 1. AePUb-HH and a polymerase pause site, 2. CMV, or 3. CMV with HH and a polymerase pause site). **C**) Predicted secondary structure of the HH ribozyme and the first five nucleotides of the DENV-2 5′UTR generated using RNAFold and visualised with VARNA. **D**) Kinetics of DENV-2 rescue via CPER using different linkers. Recovered virus, as three biological replicates, was titrated via TCID_50_ ELISA from days three to six dpt.. Error bars = SEM. LOD = Limit of Detection = 2.3 log_10_ IU/mL.

For launch of DENV-2, three conditions were assessed: AePUb-HH linker containing the polymerase II pause site in C6/36 cells; a standard CMV promoter linker in mammalian cells^16^ and a linker containing the CMV promoter followed by the HH Rbz sequence and containing the polymerase II pause site in mammalian cells. Once again, to allow for direct comparison between the various promoters, the DENV-2 genomic F1 was joined to the linker via overlap-extension PCR to account for overhang T_m_ variation in assembly. Fragments were assembled by CPER and subsequently transfected into C6/36 cells for the AePUb-HH approach, or into HEK293T cells for the CMV approach and incubated for 24 hours before co-culturing with Vero E6 cells for the CMV and CMV HH Rbz CPERs^16^.

Titration of virus harvested daily from three to six days post transfection revealed higher DENV-2 infectious titres for the constructs incorporating the AePUb-HH linker across all days in comparison to rescue with the CMV promoter in mammalian cells (Fig 5D). Upon comparison of CMV and CMV HH-launched virus titres, there was no significant difference noted for titre, nor time of rescue (Fig 5D). While these results showed a comparative advantage of the insect cell reverse genetics system for DENV-2, this result is confounded by the titration being performed on C6/36 cells, which may bias virus replication for a virus rescued in an insect cell line. Importantly, these results contradicted published data on the pause site and HH Rbz sequence for CMV, showing no improvements by their addition to the linker.

To determine whether the lack of improved CMV-mediated CPER launch efficiency following inclusion of the HH Rbz and polymerase pause sites was also evident for larger viral genomes, the comparison was extended to SARS-CoV-2, which has a substantially larger genome than DENV-2 (Fig 6A). SARS-CoV-2 genomic and linker fragments (CMV or CMV with HH ribozyme and polymerase II pause, Fig 6B, C) were amplified, purified, and assembled by CPER, which was subsequently assessed by gel electrophoresis revealing a large band at the expected size of approximately 31kb (Fig 6D). CPERs were transfected into HEK293T cells expressing ACE2 and *h*TMPRSS2 before being co-cultured with Vero E6 cells expressing *h*TMPRSS2 12 hours post-transfection. Supernatant collected daily post-transfection were titrated on Vero E6 cells expressing *h*TMPRSS2 by immunoplaque assay (iPA), where fluorescent foci were visible as early as 4 days-post transfection for the standard CMV linker launched virus and 5 days post transfection for the virus launched with the CMV linker containing HH and the polymerase pause site (Fig 6EF). However, the titres on subsequent days were relatively similar across both conditions (Fig 6F) and immunoplaque morphology was not different between the two launched viruses (Fig 6G). This result reinforced those observed with DENV-2 whereby it was demonstrated that the addition of a HH Rbz and polymerase pause site is superfluous for virus synthesis in CMV-driven mammalian systems and thus contradicting previous literature.

**Figure 6.**
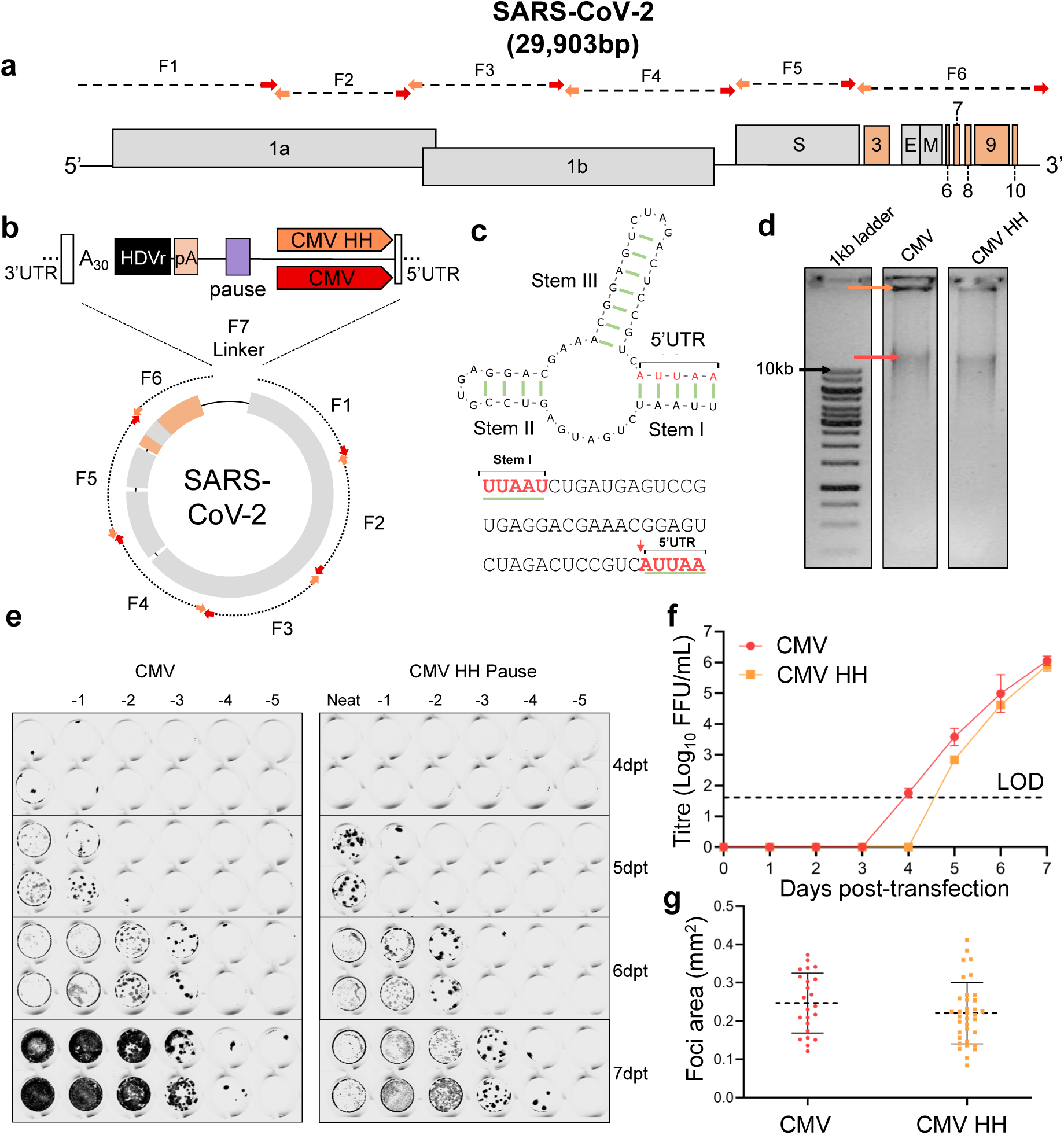
Comparison of promoter systems for SARS-CoV-2 reverse genetics. **A**) CPER schematics showing the overlapping fragments comprising the SARS-CoV-2 genome. **B**) The DNA linker was incorporated by CPER and contained the following promoters and other sequence elements: CMV alone or CMV with a polymerase pause site, and a HH Rbz sequence (CMV-HH). **C**) Predicted secondary structure of the HH ribozyme and the first five nucleotides of the SARS-CoV-2 5′UTR generated using RNAFold and visualised with VARNA. **D**) 1% agarose gel of the 2 CPERs for the CMV and CMV HH Pause conditions. Orange arrow indicates concatemeric DNA and red arrow marks the nicked circular CPER product for SARS-CoV-2. Black arrow shows the 10kb marker size for 1kb DNA ladder. **E**) iPA images of both conditions at each timepoint, from 1 to 7 days post-transfection. **F**) Transfection kinetics of the two CPER comparisons from 1 to 7 days post-transfection (n=1). FFU = foci-forming units. FFU/mL was determined by immunoplaque assay (iPA) with a C2 (anti-SARS-CoV-2 S^54^) nanoantibody. Error bars = SEM. LOD = Limit of Detection = 40 FFU/mL. **G**) Foci sizes (area in mm^2^) measured for both rescue conditions (CMV and CMV HH) of SARS-CoV-2 as determined using ImageJ. For statistical analysis between CMV and CMV HH foci sizes, unpaired t-test with Welch’s correction was used (ns = not significant). Centre horizontal bars represent the mean values and errors bars indicate the standard deviation.

## Discussion

Reverse genetics systems are foundational tools in molecular virology, enabling the study of synthetically generated viruses, the effect of mutations, and the development of recombinant virus platforms for an array of translational applications. In this study, a selection of various approaches were pursued to determine the most efficient methodological pipeline for launching positive-sense RNA viruses using bacteria-free reverse genetics systems. While there is substantial literature explaining the best practices and approaches for reverse genetics, there have been limited systematic comparisons of different standard bacteria-free reverse genetics methods, in addition to direct comparisons between promoters and the effects of transcription enhancing motifs. Through these analyses we established CPER as the optimal method for launch of long genome positive-sense RNA viruses in insect cells, ahead of Gibson and ISA. Through selection of a potent insect cell promoter and the use of a HH Rbz sequence, we further demonstrated efficient launch of nidoviruses and orthoflaviviruses in mosquito cells in as little as one day, enabling rapid recovery of otherwise difficult-to-launch constructs and scalable generation of mutant libraries. Despite the lack of improvements observed with the addition of ribozymes or polymerase pause sites reported for the mammalian-based systems^29,38^, the methodological comparisons and optimisations that we present are a useful reference point for future optimisation of those systems.

We have previously established CPER as an efficient reverse genetics system for the recovery of viruses in mammalian cells using the CMV promoter^10,16,26^ and a range of mosquito-restricted viruses, including orthoflaviviruses and alphaviruses in C6/36 mosquito cells using a truncated OpIE2 promoter^16,24,25,30,45^. However, the synthesis of larger genome viruses, such as CASV as a representative of the insect-restricted nidoviruses, regularly required 18 days of incubation to detect successful launch. We also sought to increase the recovery efficiency of difficult-to-launch mutant and chimeric viruses, which often require serial passaging for rescue, while establishing a more robust platform for generating mutant libraries, including those used for deep mutational scanning (DMS)^26^.

We selected CPER, ISA, and Gibson assembly as candidate bacteria-free methods for optimisation and comparison for CASV ZsGreen synthetic reconstruction. The results demonstrated the efficacious advantage of CPER compared to Gibson assembly and ISA when launching this large genome virus. This is at least partially attributable to the one distinctive trait of CPER, namely its single step amplification, which theoretically produces double the original DNA content^10^. In contrast, ISA and Gibson assembly stitch together the original DNA either in the reaction (for Gibson), or after transfection (for ISA). ISA was incapable of rescuing infectious virus. ISA has been used across several studies for reverse genetics of SARS-CoV-2 and orthoflaviviruses^11,47,55^, however it outsources homologous recombination assembly and ligation to the host cell. There was a less substantial difference between Gibson assembly and CPER in terms of efficiency, which is not surprising given that Gibson assembly shares a similar *in vitro* circularisation process to CPER, except that it uses an exonuclease to generate ssDNA overhangs after which junctions are sealed by a ligase. These junctions are left nicked for CPER and are instead sealed in the host cell prior to transcript production, as nicked DNA cannot serve as a template for RNA production. Despite this, the doubling of DNA content during CPER likely mitigates this ligation difference, resulting in greater yields than by the Gibson assembly method. One limitation of these comparisons herein is that their relative efficiency can vary based on the length of the overhangs and also the transfection methodology used. For instance, ISA is typically used with long overhangs upwards of 60bp^56^, while the standard overhang length used in these experiments herein was 30bp.

CPER efficiency has been enhanced recently by adding a ligation step subsequent to the circularisation step, which is enabled by the 5’ phosphorylation of PCR fragments with a T4 polynucleotide kinase prior to the circularisation^29^. Gibson Assembly has also been enhanced by phage ø29 DNA polymerase amplification of the circular product after the isothermal incubation^14^. This allows for upscaling of the final product, and therefore a higher input of transfected circular product, which may also be purified and quantified prior to transfection. Indeed, future optimisation of our CPER protocol may focus on addition of ligation or to topological modification to the DNA via supercoiling, but even through basic modification to the CPER cycle number and polymerase extension times, we were successful in reducing the time to launch CASV by at least 3 days. Furthermore, the benefit of CPER as it is applied in this paper is that it is both fast and highly efficient.

For launch of virus in C6/36 mosquito cells, comparisons of the traditional truncated OpIE2 promoter approach with the use of the *Aedes aegypti* PUb promoter, AePUb, facilitated infectious virus generation 3 days faster and to higher titres. The substantial difference in the expression capabilities was evidenced by the visual expression of ZsGreen which was inserted into the CASV genome. While OpIE2-CA showed expression of ZsGreen concomitant with virus rescue, the AePUb promoter design yielded high expression of ZsGreen prior to virus synthesis and as early as 12 hours post-transfection. Though this demonstrated how powerful the AePUb promoter is in insect cells, it does have a potential drawback in that the early signal should not be misinterpreted as being an indication of virus replication. This was best exemplified by use of the full-length AePUb linker without the addition of the HH Rbz, which showed very high ZsGreen expression from 1 day post-transfection that was not accompanied by the production of infectious virus. From these data, it was inferred that the additional nucleotides produced in the AePUb-generated transcripts are deleterious to the CASV genome, likely due to structural interference with the 5’UTR which may have roles in genome packaging and replication. Therefore, it is a necessity to include the HH Rbz appended to the AePUb sequence to maintain the authentic 5’UTR of the virus sequence intended for synthesis. This is not the case for OpIE2-CA, which was previously truncated to the transcription start site, and therefore requires no excision of extraneous nucleotides from the 5’UTR^30^. HH Rbz sites have previously been employed for T7 promoter-based mammalian reverse genetics systems successfully to prevent excess G nucleotides at the 5’end of the virus transcripts^38^.

The nuance of using a HH Rbz sequence was demonstrated for CMV in the DENV-2 and SARS-CoV-2 reverse genetics systems, both of which showed it had no beneficial impact on rescue efficacy. This is likely due to the CMV sequence having a short transcription start site relative to that for AePUb, thereby making the inclusion of a HH Rbz largely superfluous. In the absence of data on the transcription start site of a promoter sequence or data on the virus 5’UTR sensitivity to appended nucleotides, it is recommended that the HH Rbz be included for any new reverse genetics system using a novel or AePUb promoter. Overall, the results of this comparison provide a new template for a tangible improvement to mosquito virus reverse genetics and provide an optimised system for the launch of any arboviruses capable of replicating in C6/36 mosquito cells. Given the strength of this promoter in other insect cell lines^36^, there may be utility in extending its application to other reverse genetics systems in other cell lines more appropriate for the virus in question.

Previous papers have implemented a polymerase pause site derived from the human α2 globin gene to prevent polymerase readthrough, which is thought to hinder the efficiency of virus rescue via steric hindrance at the promoter site and through the generation of heterologous transcripts that do not generate viable virus^29^. This is similar to the rationale we see employed for linearising plasmids prior to *in vitro* transcription for the preparation of self-amplifying RNA or mRNA vaccines ^57^. However, there are no instances in the literature of a direct comparison of a reverse genetics system controlling for all other variables excluding the pause site. Here, a direct comparison using the AePUb promoter and the CASV ZsGreen scaffold, demonstrated no significant difference in rescue efficiency. The lack of a difference in efficiency may be due to the 5’ and 3’ ribozyme processing of transcripts via the HDVr and HH Rbz, thereby preventing generation of heterogeneous transcripts and nullifying any impact of the pause site. These results were further reinforced in the CMV HH-Rbz pause linker and CMV comparisons for DENV-2 and SARS-CoV-2, both of which showed no substantial impacts of the pause site on rescue efficiency.

The collective results of this paper represent a marked improvement to reverse genetics systems in general and for insect cells specifically. These data have facilitated the further refining of reverse genetics methodologies to isolate the specific workflows for researchers that will yield synthesis of wild-type and mutant viruses in a shorter timeframe. Specifically, the adoption of the AePUb promoter for insect cell-based reverse genetics provides a key mechanical innovation. Across both mammalian and insect reverse genetics, these data give an approximation of the optimal bacteria-free CPER-based reverse genetics system which outperforms other similar bacteria-free methods in this comparative study.

## Materials and Methods

### Cell and virus culture

C6/36 cells (*Aedes albopictus*; American Type Culture Collection CRL1660) were cultured in Roswell Park Memorial Institute medium (RPMI 1640, Gibco), with 5% fetal bovine serum (FBS) and 2 mM L-glutamine at 28°C in 5% CO_2_. Human embryonic kidney 293T cells expressing human ACE2 (HEK293T ACE2) were provided by Professor Jesse Bloom (Fred Hutchinson Cancer Research Centre, Washington, USA)^58^. African green monkey kidney cells expressing human TMPRSS2 (Vero E6-*h*TMPRSS2) were generated by transduction with lentivirus containing a puromycin-resistant human TMPRSS2 construct^16^. Mammalian cell lines were maintained at 37°C in 5% CO_2_ in Dulbecco’s Modified Eagle Medium (DMEM) supplemented with 10% FBS. Mammalian cell media was supplemented with 50 U/ml penicillin, 50 µg/ml streptomycin and 2 mM L-glutamine.

The viruses used in this study include CASV (0071 isolate; GenBank NC_023986^40^), BinJV (GenBank MG587038 ^59^), DENV-2 (ET300 isolate; GenBank MT921572), and SARS-CoV-2 (BA.5 isolate hCoV-19/Australia/QLD-QIMR03/2022 (GISAID accession ID: EPI_ISL_15671874^60^). Viruses were propagated onto Vero E6-*h*TMPRSS2 cell monolayers supplemented with 2% FBS approximately 18 hours post-seeding.

Work with reporter CASV, BinJV viruses and CPER-recovered DENV-2 was performed in a certified PC2 facility at The University of Queensland (UQ) and approved by the UQ Institutional Biosafety Committee (IBC; IBC/1289/SCMB/2020, IBC/1505/SCMB/2023; IBC/1350v2/SCMB/QIMR/2021). All infectious work with SARS-CoV-2 was performed in a certified PC3 facility at UQ under IBC approval IBC/390B/SCMB/2020, IBC1301/SCMB/2020.

### RNA extraction and cDNA synthesis

For CASV, BinJV, and DENV-2, 150µL of viral supernatant was aspirated from infected cell monolayers and inactivated using a lysis buffer (RAV1) supplied in the NucleoSpin® Viral RNA isolation column kit (Macherey-Nagel, Cat. Number: 740956.50). Viral RNA was then extracted according to the manufacturer’s protocol. For SARS-CoV-2, 250µL of concentrated viral supernatant (as per ^16^) was extracted using TRIzol™ LS (Thermo Fisher Scientific, USA) according to the manufacturer’s instruction. cDNA preparation from the viral RNA was performed using the SuperScript™ IV First-Strand Synthesis System (Thermo Fisher Scientific, USA, Cat. Number: 18091050). Gene-specific reverse primers (Table S1-9) were used for synthesis of cDNA for CASV, BinJV, and DENV-2, while random hexamers (Promega) and a dT primer were used for SARS-CoV-2. Vestiges of RNA were removed from the synthesised cDNA by RNase H treatment (New England Biolabs, Cat. Number: M0297S).

### Preparation of DNA fragments for assembly

For the CASV fluorescent ZsGreen reporter virus, fragments were amplified by PCR using PrimeSTAR GXL DNA polymerase (Takara Bio) from cDNA template (Table S1-4). The OpIE2-CA linker containing the SV40 pA site and the HDVr was PCR amplified from a plasmid template containing the entire linker sequence, as previously outlined^30^. The AePUb promoter sequence was amplified from plasmid template^37^. This sequence was joined to the linker sequence (minus the OpIE2-CA promoter) by overlap-extension PCR with GXL polymerase (Table S1). The reverse primer at the 3’ end of the AePUb promoter was designed to optionally contain the HH Rbz sequence. The poly A-tailed region containing homologous overlaps into the CASV 3’UTR and the linker was ordered as an ultramer from Integrated DNA Technologies (IDT, Singapore). It was then joined by overlap-extension PCR to the linker fragment. The ZsGreen gene was amplified by PCR from a synthetically-derived double-stranded DNA gBlock from IDT and then joined to adjacent genomic fragments using overlap-extension PCR with GXL. For the polymerase II pause site comparisons, the sequence was derived from the human α2 globin gene ^29^ and along with the full-length linker it was ordered as a synthetic gBlock from IDT and used as a template for PCR.

BinJV mCherry backbone fragments were amplified by GXL DNA polymerase from viral cDNA. The BinJV 5’UTR (96bp), the first 153bp of the BinJV capsid (C), the mCherry fluorescent protein, a foot-and-mouth disease virus (FMDV) 2A ribosome skipping peptide, the full-length codon-optimised BinJV capsid gene, and the 5′ end of BinJV prM gene were ordered as a synthetic gBlock from IDT as described previously (Table S8-9) ^45^. The cDNA was generated by SuperScript IV RT with a gene-specific 3’UTR reverse primer according to the manufacturer’s protocol (Thermo Fisher Scientific).

DENV-2 and SARS-CoV-2 genomic fragments were amplified from the cDNA template outlined above (Table S5-7, S10-11). The CMV linker was amplified from a previously generated plasmid template using GXL polymerase ^16^. The CMV pause HH Rbz linker was generated by insertion of the pause and HH Rbz sequences in the overlapping regions of primers. This involved amplification of two overlapping fragments which were then stitched using overlap-extension PCR with GXL polymerase. Following amplification of all aforementioned PCR products, the amplicons were resolved on TAE agarose gels, followed by excision of target bands and subsequent purification using a Monarch® Spin DNA Gel Extraction Kit (New England Biolabs) or a NucleoSpin® Gel and PCR Clean-up kit (Scientifix).

### Reverse Genetics Assembly Methods

Following PCR amplification and subsequent purification, amplicon concentrations were quantified using a NanoDrop One microvolume spectrophotometer (Thermo Fisher Scientific). A total of 0.1pmol of each constituent fragment was included for all reverse genetics assemblies. These were then assembled according to one of the below protocols.

#### CPER

The CPER was set up in a total volume of 50 µL as per the manufacturer’s conditions using PrimeSTAR GXL polymerase (Takara Bio). The reaction was adjusted to include 2 µL of GXL polymerase rather than the recommended 1 µL. For CPER I, the reaction was cycled at 98 °C for 2 min, followed by 20 cycles of 98 °C for 10 s, 55 °C for 15 s, 68 °C extension for 25 mins, and a final extension of 68 °C for 25 mins. For CPER II, the protocol was modified to have 35 cycles. For CPER III, the number of cycles was set to 20, the extension time was reduced to 7 minutes, and the final extension was reduced to 10 minutes.

#### Gibson Assembly

Equimolar amounts (0.1pmol) of all CASV ZsGreen fragments were included in a 20 µL NEB HiFi Assembly reaction using 10 µL of 2x NEBuilder HiFi Assembly Mix (New England Biolabs). The 20 µL reaction mix was incubated for an extended time of 4 hours at 50°C.

#### ISA

Equimolar amounts (0.1pmol) of overlapping CASV ZsGreen fragments were combined in a 0.2 µL tube immediately prior to being transferred to the TransIT-LT1 transfection complexes.

### Reverse genetics transfections

CPER, NEB HiFi Assembly were transfected into target cell lines. For insect cell experiments, C6/36 cells were seeded 20-24 hours prior to transfection to reach approximately 80% confluency. Assembly reactions or 2.5 µg of plasmids were combined with 7.5 µL of TransIT-LT1 (Mirus Bio) and 250 µL of OptiMEM Reduced Serum medium (Gibco) and incubated at room temperature for 30 minutes before being added dropwise onto cells containing fresh RPMI 1640 media supplemented with 2% FBS.

For DENV-2 mammalian cell transfections, HEK293T cells were seeded for 80% confluency in a 6 well plate 20 hours prior to transfection. CPER assemblies were then incubated for 30 minutes with 7.5 µL of TransIT-2020 (Mirus Bio) and 250 µL of OptiMEM Reduced Serum medium. At 24 hours post-transfection, transfected cell monolayers were trypsinized and co-cultured with 70% confluent Vero E6 cells in DMEM supplemented with 2% FBS.

For SARS-CoV-2 transfections, HEK293T-ACE2-TMPRSS2 cells were seeded for 80% confluency in a 6 well plate 20 hours prior to transfection. Full CPER mixes were incubated with Lipofectamine™ LTX Reagent with PLUS™ Reagent (Invitrogen, Thermo Fisher Scientific) and OptiMEM Reduced Serum medium and transfected by addition to cell monolayers according to the manufacturer’s protocol. At 12 hours post-transfection, transfected cell monolayers were trypsinized and co-cultured with 70% confluent Vero E6-*h*TMPRSS2 cells in DMEM supplemented with 10% FBS.

### Rescue kinetics

At each timepoint post-transfection, 300 µL of media was harvested from transfected cells and stored at −80°C. For CASV ZsGreen, BinJV mCherry, and DENV-2, the harvested supernatant was titrated onto C6/36 cells according to a TCID_50_ protocol described in previous studies^61^. Briefly, viral supernatant was diluted 1 in 10 in RPMI containing 2% FBS and added in quadruplicates to the first column of a 96 well plate. Virus was titrated across 11 columns in 10-fold dilutions in cell culture media before addition to cell monolayers. CASV ZsGreen and BinJV mCherry were scored by the presence of fluorescence using a ZOE Fluorescent Cell Imager (Bio-Rad). For DENV-2, virus-positive wells were determined by fixed-cell ELISA, as previously detailed^24^, but with fixation of the cell monolayers fixed in a solution of 4% formaldehyde and 0.5% Triton X-100 in PBS. Briefly, these were then incubated with a blocking buffer solution (0.05 M Tris-HCl [pH 8.0], 1 mM EDTA, 0.15 M NaCl, 0.2% casein, and 0.05% Tween 20). After blocking, the plates were probed with anti-orthoflavivirus NS1 mAb 4G4. Plates were then washed with PBS containing 0.05% Tween 20 (PBST). Subsequently, goat anti-mouse Ig conjugated to horseradish peroxidase (HRP) (P0447, Dako) was added and incubated for 1 hour before washing with PBST. Lastly, a substrate solution comprising 1 mM 2,2-azinobis(3-ethylbenzthiazoline-6-sulfonic acid) and 3 mM H_2_O_2_ in a buffer prepared by mixing 0.1 M citric acid with 0.2 M Na_2_HPO_4_ to give a pH of 4.2 was added before incubation at room temperature for 1 hour. Absorbance was measured at 405 nm, and each well was identified as positive for OD values that were greater than 2x the average of the mock uninfected cell monolayers. For SARS-CoV-2, supernatant was harvested at each timepoint and titrated onto Vero E6-*h*TMPRSS2 cells for assessment by iPA (see method below).

### Immunoplaque Assay

Immunoplaque assays were conducted as previously described^62^. Briefly, 5 × 10^4^ Vero E6-hTMPRSS2 cells were seeded per well in 96-well plates. Samples were serially diluted 10-fold in DMEM supplemented with 2% FBS and P/S, and 25 μL of each dilution was added to the cell monolayer. Following a 1 hr incubation, 175 μL of overlay medium was added and the plates were incubated for a further 24 h. The overlay was then removed, and the plates were fixed for 1 h with ice-cold 80% acetone in PBS before being air-dried completely. Plates were blocked with 5% KPL blocking solution prepared in PBS-T for 1 h at room temperature. For primary incubation, the human Fc-linked anti-SARS-CoV-2 nucleocapsid nanobody C2^63^, kindly provided by Dr Ariel Isaacs (School of Chemistry and Molecular Biosciences, The University of Queensland), was added at 1 μg/mL and incubated overnight at 4°C, followed by three washes with PBS containing 0.05% Tween 20 (PBS-T). Plates were then incubated with goat anti-human IRDye 800CW secondary antibody (LI-COR, USA) at 1 μg/mL in blocking solution for 1 h at 37°C, followed by three washes with PBS-T. Dried plates were scanned using an Odyssey CLx Imaging System (42 μm resolution, medium quality, 3.0 mm offset). Foci-forming units (FFU) were quantified using Image Studio Lite (v5.2.5; LI-COR, Lincoln, NE, USA), and virus titres were calculated as FFU/mL. Immunoplaque area (mm²) was measured using ImageJ (version 1.54f; National Institutes of Health, Bethesda, MD, USA).

### Lateral Flow Assay

Purified anti-CASV mAb 5D3^48^ was conjugated to 40 nm carboxyl gold nanoparticles (NanoComposix, San Diego, CA, USA) as per manufacturer’s instructions. The conjugated gold was applied to pre-blocked conjugate pads (STD17, 370 um, Cytiva) as previously described^64^. Pre-laminated nitrocellulose membranes (Cytiva) were striped with purified anti-CASV mAb 9D7^48^ at 1 mg/ml as the capture mAb and goat anti-mouse IgG (Invitrogen) as the control at 0.9 μl/cm. Assembled strips were assessed with 40 μl cell culture supernatant and allowed to develop for 15 minutes. Strips were read using a LeeLu colorimetric reader (Lumos) and peak above background recorded and graphed as previously described^64^.

## Acknowledgements

This project was funded by an Advance Queensland Industry Research Fellowship awarded to J H-P (AQIRF067-2020-CV). J.R.P was supported by a Research Training Program Stipend from the University of Queensland. The SARS-CoV-2 component of this study was supported by an NHMRC Ideas Grant (2012883) awarded to A.A.K. A.K. was supported by the UK Medical Research Council (MC_UU_12014). We thank Dr Ariel Isaacs for generously providing the C2 anti-SARS-CoV-2 nucleocapsid nanobody used in this study. We thank Dr Jessica Harrison, Humda Zainab, and Dylan Bowman for their excellent technical assistance, and Professors Gorben Pijlman, Jeroen Kortekaas and Jelke Fros (Wageningen University and Research, Netherlands) for insightful discussions. Finally, we acknowledge BioCifer Pty Ltd (Queensland, Australia) for the use of LFT manufacturing equipment.

## Supplementary Files

**Table S1.**
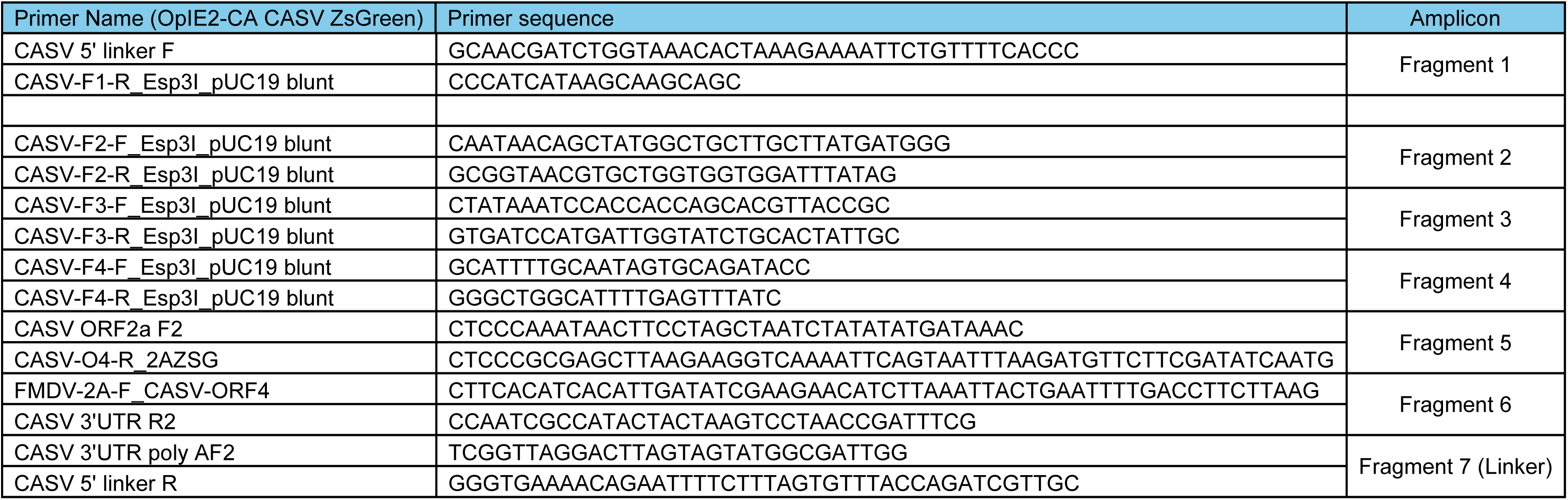
CASV OpIE2-CA construct primers for fragments.

**Table S2.**
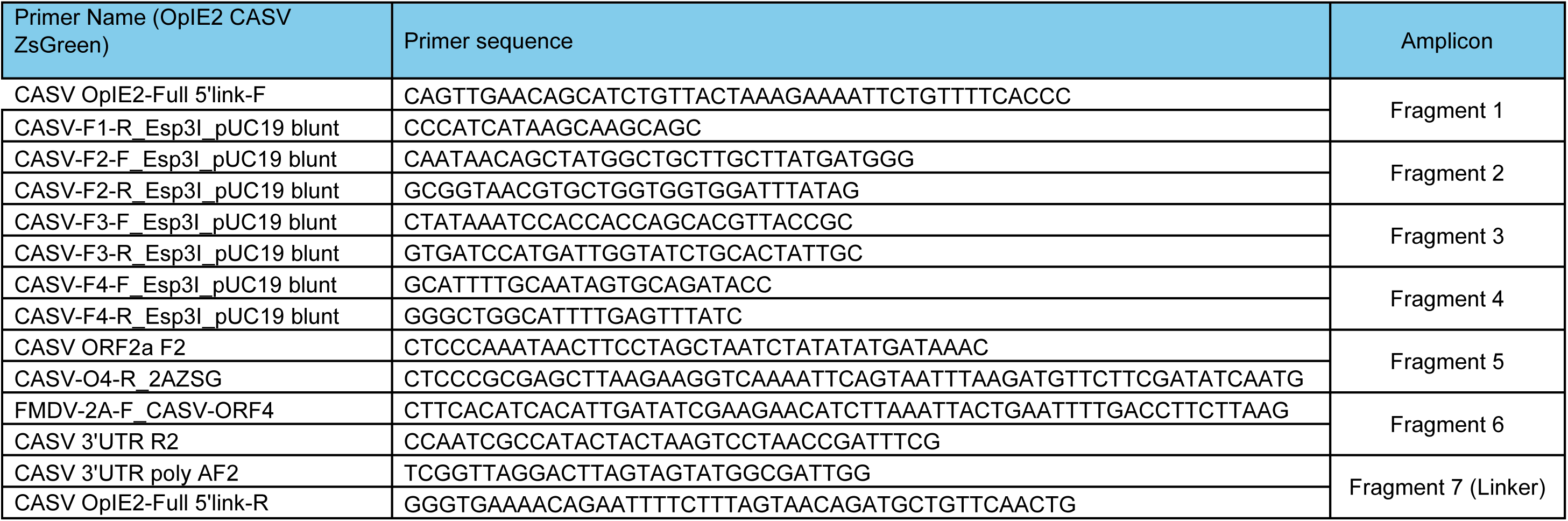
CASV OpIE2 construct primers for fragments.

**Table S3.**
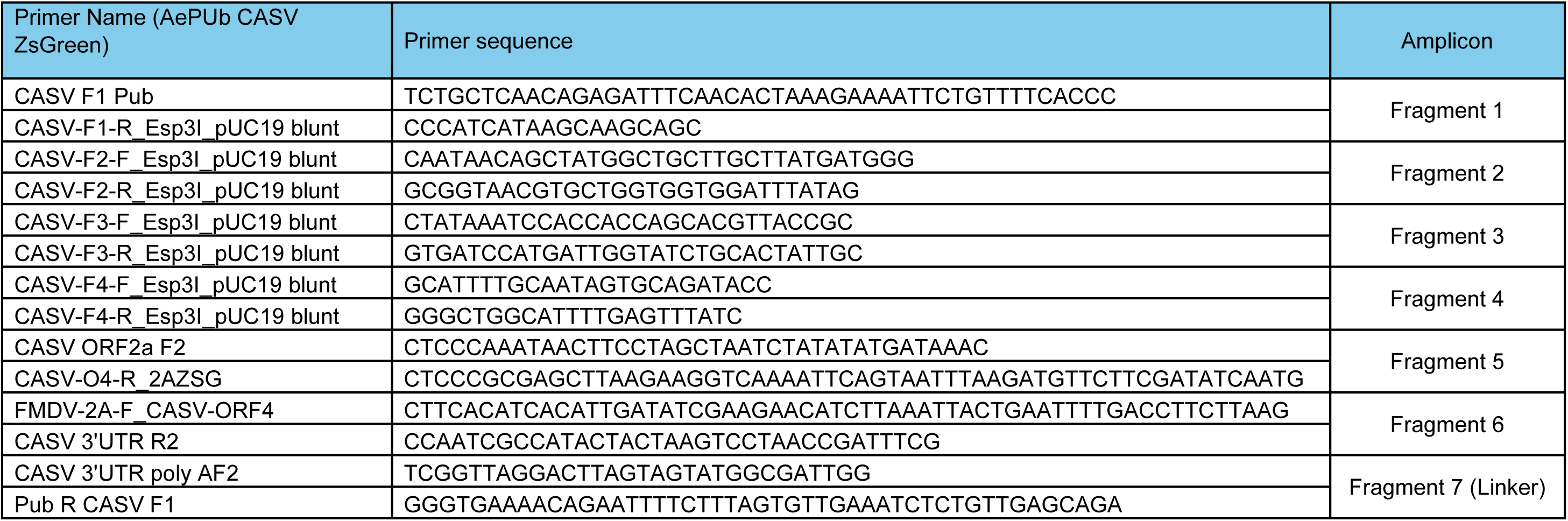
CASV AePUb construct primers for fragments.

**Table S4.**
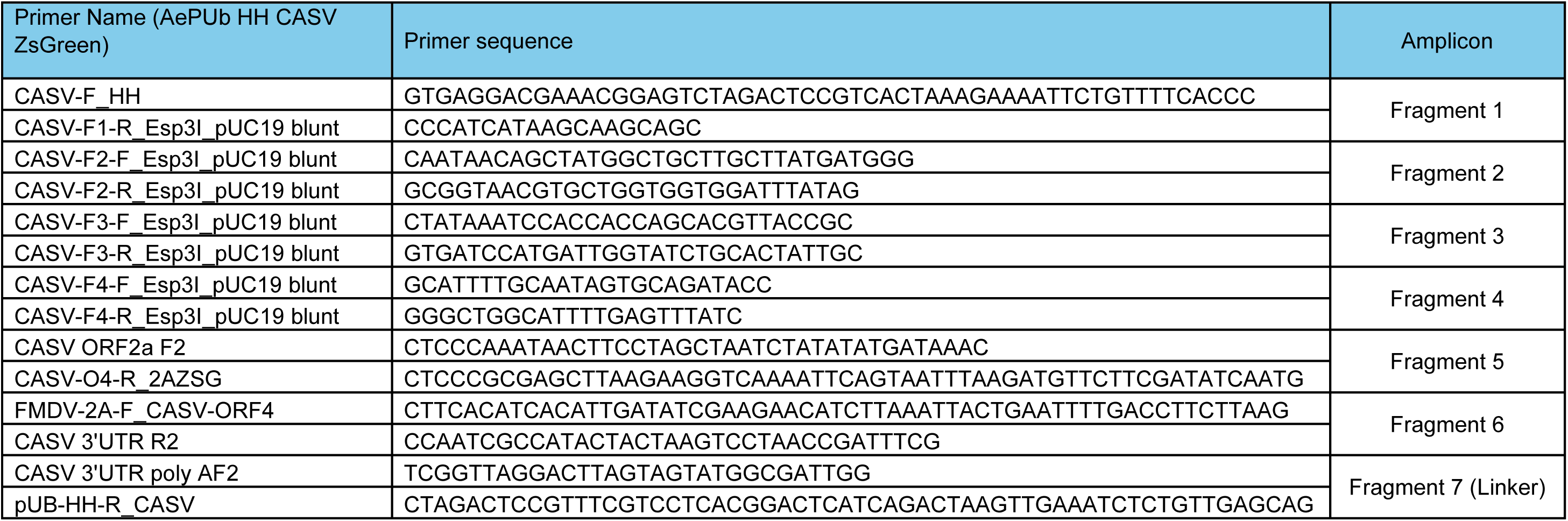
CASV AePUb HH construct primers for fragments.

**Table S5.**
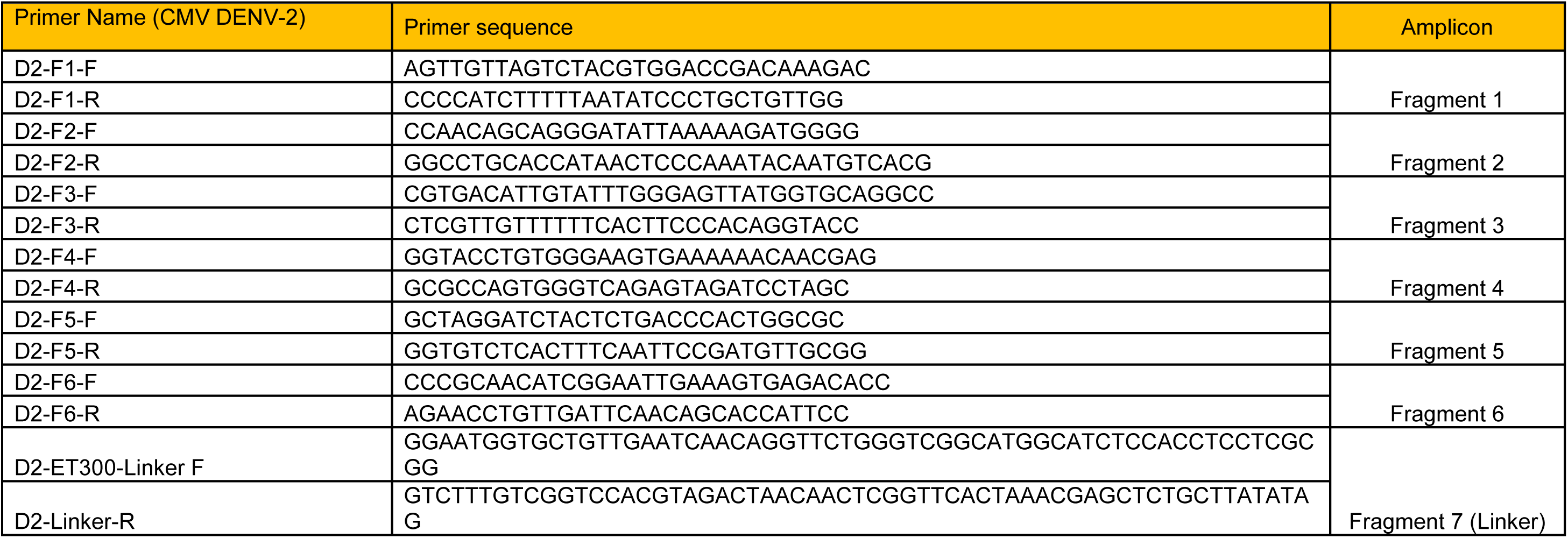
DENV-2 CMV construct primers for fragments.

**Table S6.**
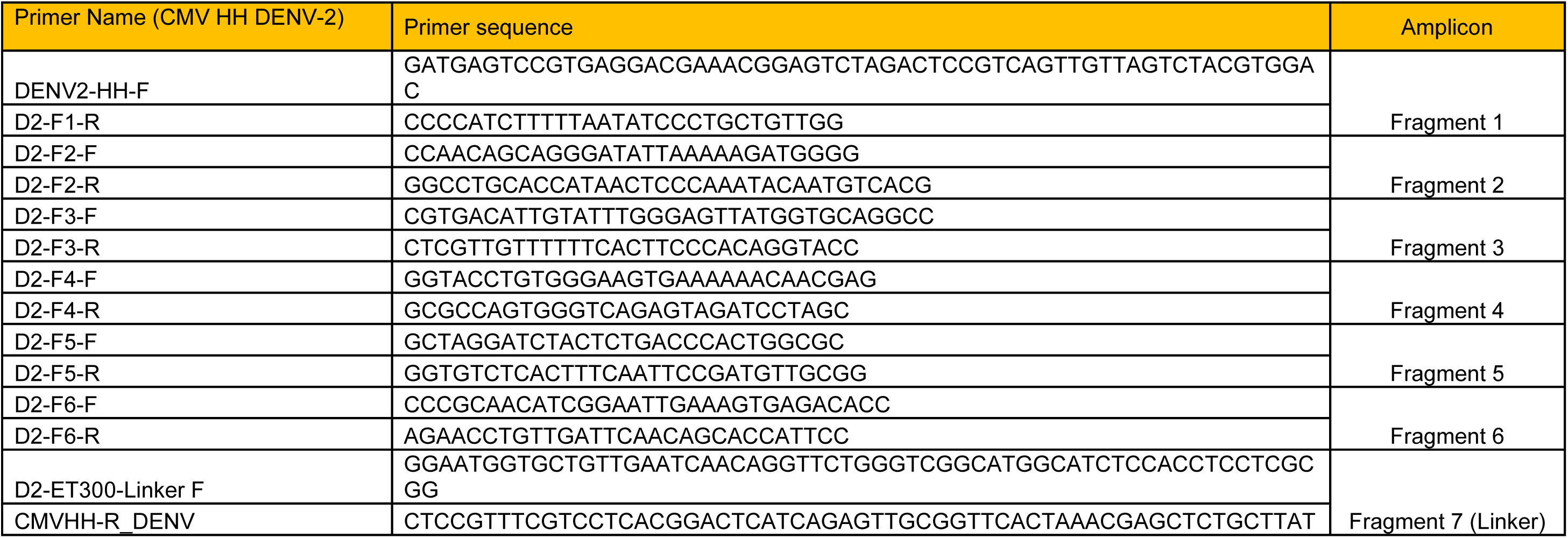
DENV-2 CMV HH construct primers for fragments.

**Table S7.**
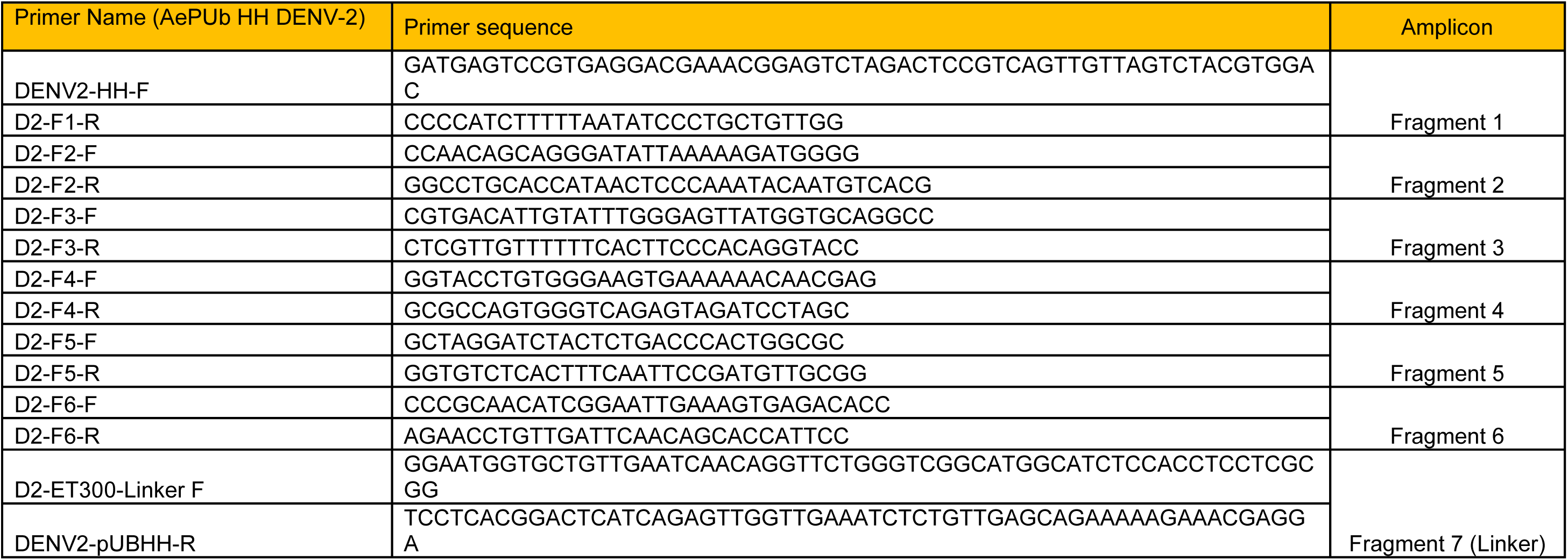
DENV-2 AePUb HH construct primers for fragments.

**Table S8.**
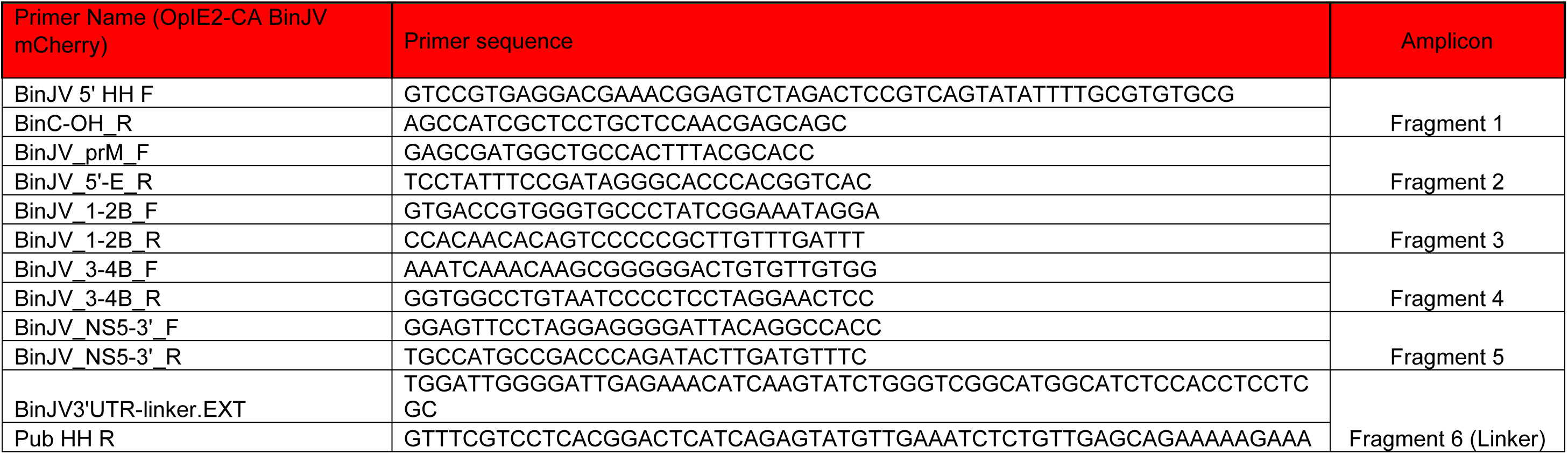
BinJV mCherry OpIE2-CA construct primers for fragments.

**Table S9.**
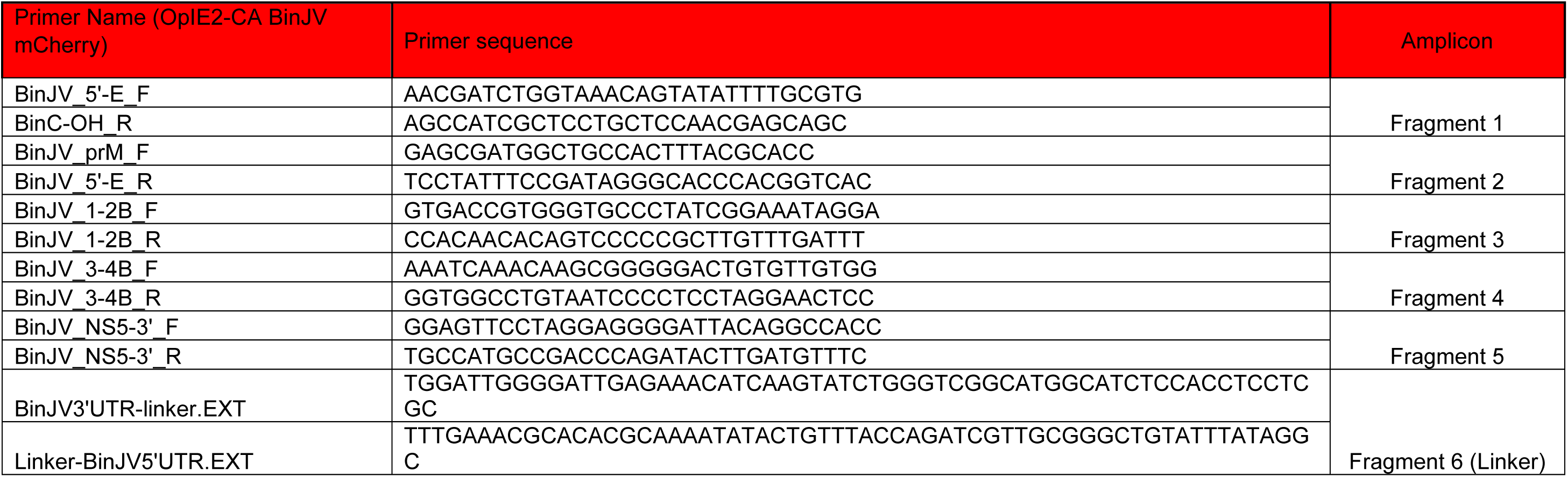
BinJV mCherry AePUb construct primers for fragments.

**Table S10.**
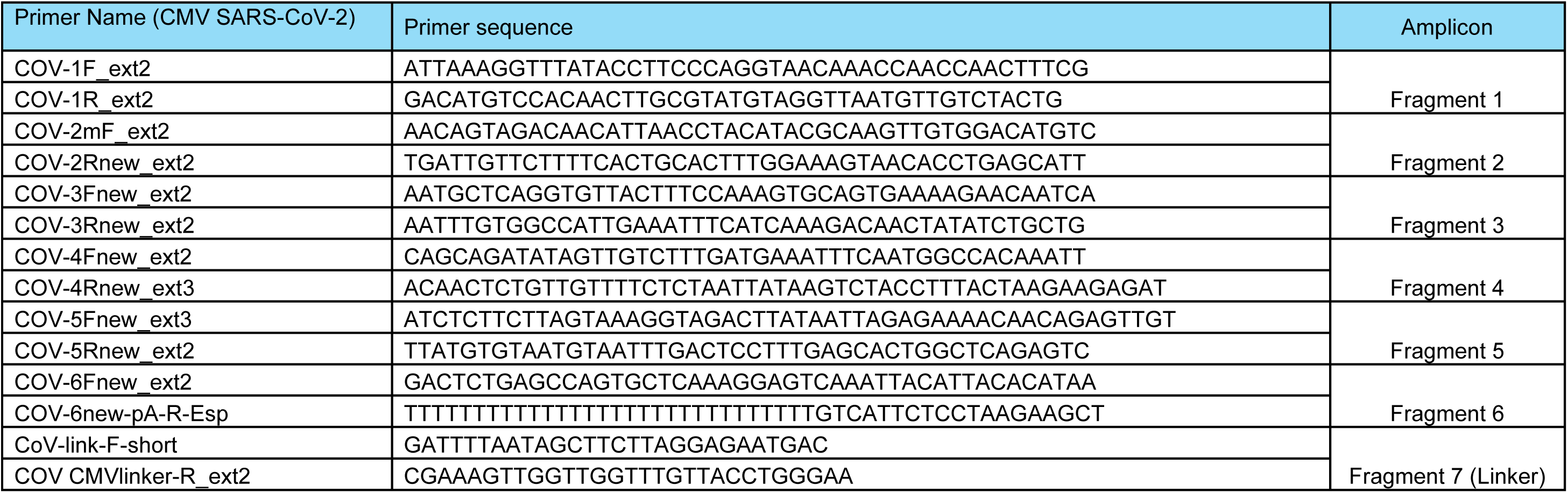
SARS CMV construct primers for fragments.

**Table S11.**
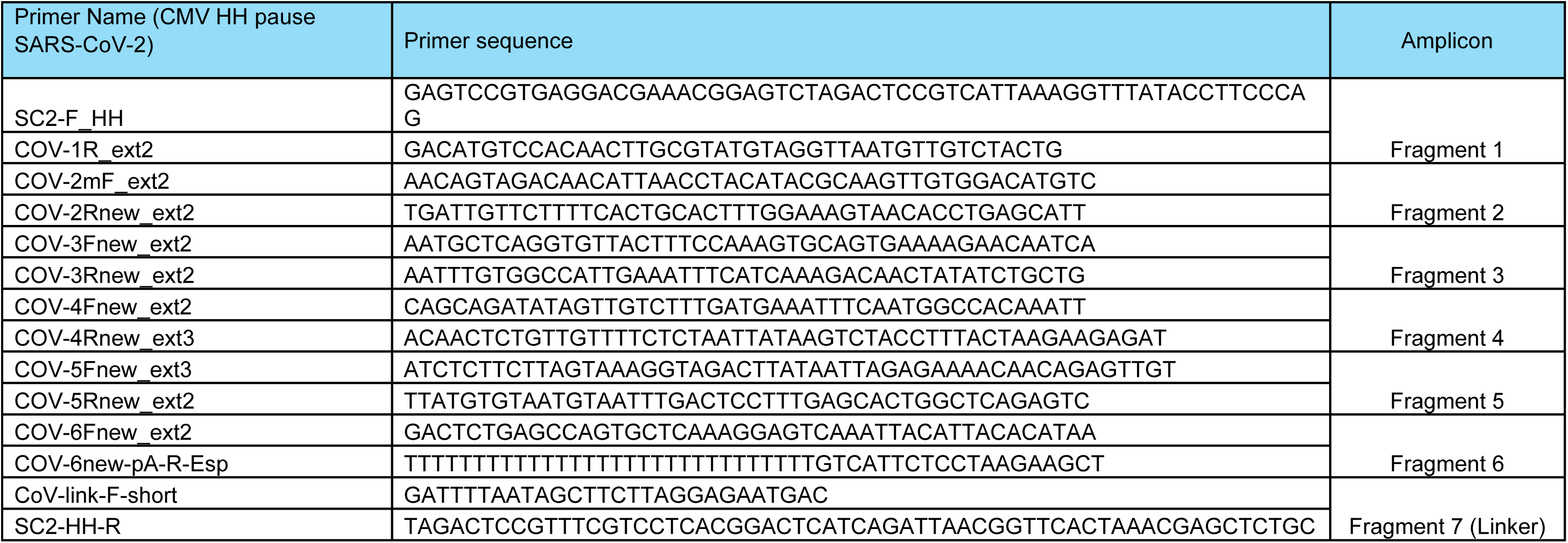
SARS CMV HH construct primers for fragments.

